# *SlHSFB3a* developmentally regulates lateral root formation by modulating auxin signaling in tomato

**DOI:** 10.1101/2025.04.15.648910

**Authors:** Adity Majee, Babythoithoi Sairem, Vinod Kumar, Aniruddha P. Sane, Vidhu A. Sane

## Abstract

- The function of HSFs, known otherwise as ‘master thermoregulators,’ in plant developmental remains largely uninvestigated.
- In this study, we strategically analyze *SlHSFB3a*, a class B sub member of HSF transcription factor family, uniquely expresses in age-dependent tomato roots and improves root architecture by synchronizing auxin homeostasis.
- This data demonstrates *SlHSFB3a* overexpressed transgenics display higher lateral root (LR) density and early LR emergence improving root architecture. Generation of CRISPR-Knockout mutants displayed contrasting phenotype, confirming *SlHSFB3a*’s vital role in root growth. In *SlHSFB3a* manipulated roots, concentration gradient auxin responses oscillated with increase in LR number.
- We highlight the signal transduction of *SlHSFB3a* mediated auxin activation that enhances tomato LRs. *SlHSFB3a* directly inhibits auxin repressors, increases auxin flow via ARF7/LOB20 pathway and positively modulates LR growth.

## Introduction

Plants are inherently dynamic systems, constantly adapting to bolster their ability to withstand and adjust to ever-changing environmental conditions. This adaptability is orchestrated by intricate metabolic networks and signaling pathways, finely regulated by genetic and hormonal mechanisms that enable plants to respond to diverse environmental cues. Central to this adaptive process is the root, a fundamental organ that anchors the plant, provides structural support, and enables nutrient and water uptake critical for growth. Roots are typically the first to encounter and react to adverse environmental conditions, requiring rapid and resilient adaptations (Siddiqui *et al*., 2021). The genetic basis for stress response is governed by transcription factors that mediate phytohormonal signaling, acting as molecular switches to fine-tune gene expression(Wagh *et al*., 2023).

Among these transcription factors, heat shock transcription factors (HSFs) serve as central regulators of heat responses, including protein biosynthesis, folding, and metabolic changes. During terrestrial adaptation, HSFs have expanded their roles, encompassing growth responses in addition to their conventional functions (Lehti-Shiu *et al*., 2017; Gonzalez-Bayon *et al*., 2019). Traditionally viewed as thermoregulators, recent findings reveal that HSFA1 also regulates antheridophore development in *Marchantia polymorpha* under 22°C non-heat stress conditions, likely through auxin accumulation (Wu *et al*., 2022). Tissue and organ specific transcriptome in *Marchantia* identified co-expression of cell wall and development-related genes, emphasizing HSFB1’s role in altered morphology under non-heat conditions (Wu *et al*., 2022). Peach *HSF5* transformation in *Arabidopsis* resulted in dwarf and deformed primary roots (Tan *et al*., 2021), while ectopic expression of *AtHSFB4* reduce root length and alter root meristem morphology (Begum, Reuter & Schöffl, 2013). In tomato, mutated HSFB4a (an ortholog of *Arabidopsis SCHIZORIZA*) caused bushy root phenotype with shorter primary roots, linked to a serine-to-cysteine substitution in the DNA-binding domain of *SlHSFB4a* (Kevei *et al*., 2022).

Tomato (*Solanum lycopersicum*), a major crop consumed worldwide, emerged as a model for studies on fruit development and biotic and abiotic stress (Feder *et al*., 2020). However, its root research remains limited due to the challenges posed by its underground structure, which hinders direct observation of root physiology (Böhm, 2012). Auxin is the key hormone coordinating primary and lateral root growth in tomato, with transcriptomic studies identifying differentially regulated auxin-related genes, including *AUX/IAA, ARF, SAUR*, and *SlGH3*, in young roots (Gupta *et al*., 2013; Kumar *et al*., 2021). 80 Cytokinin-related genes found to be differentially regulated in tomato PR and LR tissues, notably *SlCKX*s and Type A Response Regulator (*SlRRs)*, suggesting cytokinin-mediated root growth signaling (Gupta *et al*., 2013). Other phytohormones namely, ABA, ethylene and strigolactone are reported to form interaction networks in tomato that synchronizes root growth, but further in-depth studies regarding the molecular mechanism is yet to be unraveled. This study focuses on understanding tomato root through the combined lens of physiology and genetic crosstalk underlying root traits, that improves crop productivity and quality.

Our group’s previous transcriptomic data has shown elevated HSF transcripts in tomato root tissues under non-stress conditions, with significant shifts upon hormonal treatment (Majee *et al*., 2023). Specifically, *SlHSFA7* and *SlHSFB3a* were predominantly expressed in roots. This study investigates the role of *SlHSFB3a* and its downstream targets in regulating the signal transduction pathways critical for tomato root growth.

## Materials and methods

### Plant materials and growth parameters

Tomato seeds of Ailsa Craig variety were obtained as a gift from Prof. JJ Giovannoni (University of Cornell, Ithaca, USA). Tomato plants were grown in glasshouse, under specific conditions of 23±2°C with 16h/8h light and dark photoperiod at 100 μE m^−2^s^−1^. Root and leaf (15, 30 and 60 days old), stem, flower, seeds (green and red mature fruit) and pulp of mature fruit (green and red) were sampled by freezing in liquid nitrogen, stored in −80°C for future use. All tomato transgenic lines were grown in glasshouse maintaining at 23±2°C with a 12 h light/dark photoperiod at 100 μE m^−2^s^−1^.

### Sequence Analysis, phylogenetic analysis

Tomato genome files were downloaded from phytozome v9 portal (phytozome.jgi.doe.gov/pz/portal.html). The SlHSFB3a sequence was analysed for conserved motifs in NCBI-CDD (https://www.ncbi.nlm.nih.gov/Structure/cdd/wrpsb.cgi). For multiple sequence alignment, the amino acid sequences of 14 closest orthologues of SlHSFB3a and an outgroup from *Saccharomyces cerevisiae* were aligned by clustal omega and the conserved domains were searched in CDD, represented via shady box domain. Next, to understand the phylogenetic relationship, an unrooted tree was constructed using maximum likelihood algorithm with 2000 bootstrap replicates.

### Repressor Analysis

The SlHSFB3a full-length coding sequence was cloned into the bait vector pGBKT7 using tomato cDNA as a template. pGBKT7:SlHSFB3a and pGADT7 were co-transformed into yeast (*Saccharomyces cerevisiae*) strain AH109 according to Clontech Yeast Protocols Handbook (TakaraBio, USA). Similarly, the positive (pGBKT7-53 co-transformed with pGADT7-T) and negative (empty pGBKT7) controls were transformed into yeast. The resulting positive colonies were grown on synthetic defined (SD) medium lacking leucine and tryptophan (SD/-L-W) for 3 days at 30°C. Then, the confirmed positive colonies were flooded with X-gal buffer, incubated at 37°C till blue color appears.

For SlHSFB3a *in-planta* repressor analysis, *Arabidopsis* protoplasts were extracted and transformed with effector (GAL4DBD), GAL4DBD-*SlHSFB3a* and reporter [GAL4(4X)::GUS] constructs following the protocol described in (Kumar *et al*., 2024). β-glucuronidase activity was quantified after 20h incubation.

### Generation of *SlHSFB3a* over-expression and knockout transgenic lines in tomato

For *SlHSFB3a* functional characterization, over-expression (OEx) and knockout (KO) constructs were prepared. For OEx construct, *SlHSFB3a* ORF was cloned in sense orientation driven under CaMV35S promoter in pBI121.The pBI121:SlHSFB3a construct was transformed to tomato cotyledons via agrobacterium mediated transformation. For transgene confirmation, genomic DNA isolated from individual transgenic lines was used as a template for genotyping using vector and gene specific primers listed (Supplementary Table 1).

For the KO lines, a 20bp guide RNA was designed (CHOPCHOP; chopchop.cbu.uib.no) and cloned into pHSE401 vector with *Cas9* endonuclease and transformed to generate transgenic lines. T0 plants were screened for mutation detection via mismatch cleavage assay following the protocol described in Guide-it^TM^ Mutation Detection Kit Protocol-At-A-Glance (TakaraBio, USA). Mutations in *SlHSFB3a* CDS was confirmed by Sanger sequencing (Eurofins, India)

### Root phenotypic analysis

WT, *SlHSFB3a* OEx and KO seeds were sown in 1/2 MS agar media and morphological patterns were observed with respect to primary and lateral root growth. The quantitative analysis of the root structural parameters calculated using ImageJ. For, LR initiation 4 DAG tomato seedlings were documented by Leica EZ4E (Leica Microsystems, Germany). Water displacement method is employed to measure root volume (Harrington, Mexal & Fisher, 1994).

### Tissue collection, RNA extraction and qRT-PCR

Age dependent tomato tissues were used as a template for RNA isolation from three independent biological replicates. Ground tissues were subjected for total RNA extraction using FavorPrep Plant Total RNA Mini Kit (Favorgen biotech corp, India). The RNA samples were quantified by Nanodrop (NanoDrop OneC, Thermo Scientific, US). For cDNA synthesis, reverse transcriptase enzyme (Puregene, Genetix Biotech Pvt. Ltd, India) was used as per instructions provided. qPCR was performed using SYBRgreen (Puregene, Genetix Biotech Pvt. Ltd, India) and ABI Step One Plus Real-time PCR machine (Applied Biosystems Inc, USA). For differential expression analysis, 2^−ΔCt^ was calculated (Ct of target − Ct of reference) and 2^−ΔΔCt^ was calculated (ΔCt mutant − ΔCt WT). *SlCAC* were used as reference gene. Primers used are listed in Supplementary Table 1. For qPCR, all samples were taken in biological and technical replicates.

### Transcriptome sequencing

Total RNA from the root tissues of WT, *SlHSFB3a* Ox and KO (30 DAG) were extracted in biological replicates and sequenced in Illumina Novaseq sequencing platform at Novelgene Technologies (Hyderabad, India). Excluding the low-quality reads, the filtered reads were processed based on quality score Q<20 parameter, mapped against *Solanum lycopersicum* SL3.0.54 genome downloaded from EMBL database (www.ebi.ac.uk). Heatmaps and volcano plots prepared using pheatmap and VolcanoseR (huygens.science.uva.nl/VolcaNoseR/). VENN diagram with (https://bioinformatics.psb.ugent.be/webtools/Venn/) and GO analysis with The Gene Ontology Resource (https://geneontology.org).

### Yeast one Hybrid

For protein-DNA interactions, bait/reporter sequences, Pro*SlIAA2* - pAbAi and Pro*SlIAA3* - pAbAi constructs were prepared using respective primer sets and transformed in Y1H Gold yeast strain following the manufacturer’s protocol (Takara, Japan). The transformed colonies were selected on SD/-Ura agar plates containing Aureobasidin-A (AbA) optimized at 100 ng ml^−1^ and incubated at 30ºC for 3-5 days. For prey construct, SlHSFB3a (ORF) cloned in pGADT7 vector was considered. Yeast competent cells of bait constructs (that were obtained from transformed positive colonies grown on SD/-Ura) were prepared and the prey construct was transformed into these yeast competent cells and selected on SD/-Leu/AbA (100ng/mL). These yeast recombinants (bait-prey constructs) were allowed to grow on SD/-Leu/AbA (100 ng/mL) at 30ºC for 3-5 days. pAbAi:p53 and mutated pAbAi vector with p53 were used as positive and negative controls.

### Dual Luciferase Assay

*SlIAA2* and *SlIAA3* promoters (~1.9 Kb, ~1.6 Kb) were used to drive luciferase expression in tobacco leaves by cloning in the gateway cloning vector, pGWB435 (Invitrogen). *SlHSFB3a* CDS was separately cloned in pGWB402 to prepare an effector construct. Empty vectors (pGWB435, pGWB402) and Pro:Luc reporters were used as negative and positive control. The constructs were agro-infiltrated (*A. tumefaciens* strain GV3101). The agro-induction media was prepared from 10mM MgCl_2_, 10mM MES (pH-5.6), 100µM acetosyringone and 0.5% glucose. Culture of the bait and prey constructs under the interaction study was mixed in 1:1 ratio of OD=1 each followed by syringe infiltration into expanded young leaves of *N. benthamiana* and then kept in dark for 24h followed by light of 12-18h. 1mM D-luciferin was evenly spread on the leaf surface and then visualized under Chemi-Doc (ChemiDoc TM XRS+ imaging system) (Zhou et al., 2018).

### Statistical Analysis

For the repressor analysis, the homogeneity of variances was calculated using the variance equality rule. After confirming that the population variances were equal a student’s t-test was conducted in MS Excel using the “t-test: Two-Sample Assuming Equal Variances” data analysis tool. For the rest of data, comparisons between data sets were performed using one-way ANOVA, followed by mean separation with Duncan’s Multiple Range Test (DMRT) as a post-hoc test. The statistically analyzed data was graphically represented as mean±SE by GraphPad prism software.

## Results

### *SlHSFB3a*: A root-localized HSF family transcriptional repressor in tomato, induced by auxin, cytokinin, and ABA

*Two HSFs i*.*e*., *SlHSFA7* and *SlHSFB3a* exhibited higher transcript levels in tomato roots as compared to other tissues under unstressed conditions (Majee *et al*., 2023). *In silico* analysis identified a 714 bp coding region for *SlHSFB3a* (Solyc04g016000), encoding a protein of approximately 27 kDa. Conserved domain analysis revealed a 94-aa DNA-binding domain (Supplementary Fig. **1a**), along with an oligomerization domain, nuclear localization sequence (NLS), and an N-terminal repressor motif (M/LFGV). Phylogenetic analysis, comparing the SlHSFB3a sequence with its other plant orthologues, indicated that SlHSFB3a is closely related to *Solanum pennellii* HSFB3 (SpHSFB3) and *Solanum tuberosum* HSFB3 (StHSFB3) (Supplementary Fig. **1b**). Plant HSFs are categorized into three classes A, B and C. Class B HSFs contain an LFGV repressor motif, characterizing the transcriptional properties of this plant-specific HSF group.

To examine *SlHSFB3a* tissue-specific expression, qRT-PCR was performed across different tomato tissues, including age-dependent roots and leaves, stem, flowers, fruit pulp, and seeds. *SlHSFB3a* was predominantly expressed in roots at 15, 30, and 60 days after germination (DAG), with maximum expression at 60 DAG, suggesting its involvement in root growth (Fig. **1a**).

**Fig. 1:**
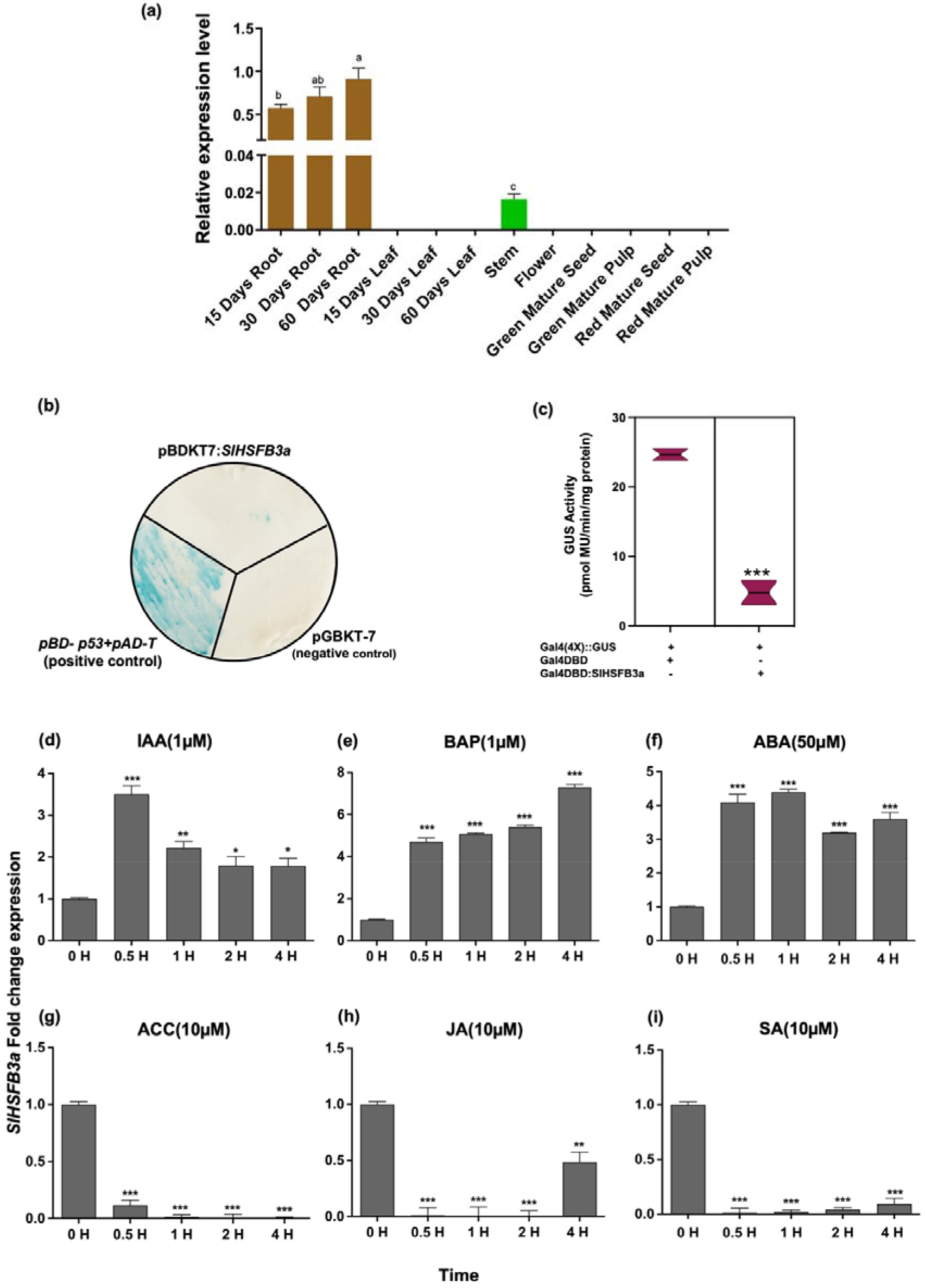
*SlHSFB3a* functional relevance in tomato (Ailsa Craig) tissues (A) qRT-PCR validation of *SlHSFB3a* transcript abundance in different tissues. *SlCAC* was used as an internal reference. Significant variation is represented by a, b and c letters and analysis by DMRT (Duncan Multiple Range Test). (B) Demonstration of *SlHSFB3a* transactivation properties. (C) Relative GUS activity of full-length (BD-SlHSFB3a) and {GAL4(4X)::GUS (reporter construct)} as determined by MUG assay. (D) *SlHSFB3a* phytohormone profiling upon hormone treatments. WT seedlings were submerged in different hormones including (D) 1μM IAA (E) 1μM BAP (F) 50 μM ABA (G) 10μM ACC (H) 10 μM ACC and (I) 10μM SA. Y-axis depicts 2^−ddCT^ values. Error bars represent the standard error (±) calculated from three independent biological replicates. Significant variation is indicated by asterisks *P < 0.05, **P < 0.01, and ***P <0.001 (single factor ANOVA).

A yeast transactivation assay confirmed that SlHSFB3a does not function as a transcriptional activator, as yeast cells transformed with a GAL4DBD-SlHSFB3a fusion did not display blue coloration, unlike the positive control (Fig. **1b**). Similarly, GAL4DBD-SlHSFB3a reduced GUS activity compared to the control, in *Arabidopsis* protoplasts, confirmed repressor nature of SlHSFB3a (Fig. **1c**).

The expression of *SlHSFB3a* was significantly induced upon treatment with 1 μM IAA, 1 μM BAP, and 50 μM ABA at various time intervals (0, 0.5, 1, 2, and 4 hours), as illustrated in Fig. **1(d-i)**. Within 30 minutes of IAA exposure, *SlHSFB3a* transcripts increased 3.5-fold compared to untreated. Although the expression gradually declined with prolonged exposure, it remained higher than the untreated even after 4 hours (Fig. **1d**). Similarly, 1 μM BAP elevated *SlHSFB3a* expression within initial 30 minutes, reaching a maximum 6-fold increase after 4 hours. Treatment with 50 μM ABA also upregulated *SlHSFB3a* expression. In contrast, exposure to 10 μM JA, 10 μM SA, or 10 μM ACC rapidly suppressed *SlHSFB3a* expression {Fig. **1(e-i)**}.

### *SlHSFB3a* promotes lateral root development in age-dependent tomato plants

Overexpression (OEx) and knockout (KO) *SlHSFB3a* tomato lines (Supplementary Fig. **2b**) showed significant differences in transcript levels. Sanger sequencing revealed a single insertion mutation (+T), and an additional G to A substitution in *SlHSFB3a* KO L-8. The insertion mutation led to a premature stop codon in the SlHSFB3a protein sequence (Fig. **2a**). In two independent CRISPR-edited lines, *SlHSFB3a* transcripts were undetectable, confirming them as *SlHSFB3a* knockout (KO) lines (Fig. **2b**). In the root tissues of 15 DAG OEx transgenics, *SlHSFB3a* transcripts were enhanced by approximately 3-5-fold (Fig. **2b**). Phenotypic analysis of the transgenic roots revealed notable changes in tomato root architecture. In *SlHSFB3a* OEx (15 DAG), primary root (PR) length was shorter compared to wild-type (WT) and KO (Fig. **3 a,b**). PR length averaged 6.07 ± 0.73 cm, 6.73 ± 0.65 cm, and 6.95 ± 0.25 cm across three OEx lines, while WT plants averaged at 9.54 ± 1.7 cm. Lateral root (LR) number increased substantially in OEx, whereas KO exhibited 33-34% reduction (Fig. **3c**). LR density, a measure of LR spacing along PR length, was 1.7-2-fold higher in OEx roots and lower in KO (Fig. **3d**). Increased LR density, indicates that the formation of lateral root primordia (LRP) as a potential contributor of this phenotype. In germinated seeds, LR emergence was observed over 8 days. OEx seedlings initiated early LRP emergence on day 4, while KO displayed delayed LR formation, with buds appearing on day 6 (Fig. 3e). Stereo microscopic images of 4-DAG seedlings further supported this, showing elongated LRPs in OEx roots that were absent in WT and KO (Fig. **3f**). The findings confirm that *SlHSFB3a* promotes LRP formation, accounting for the increase in LR number and density.

**Fig. 2:**
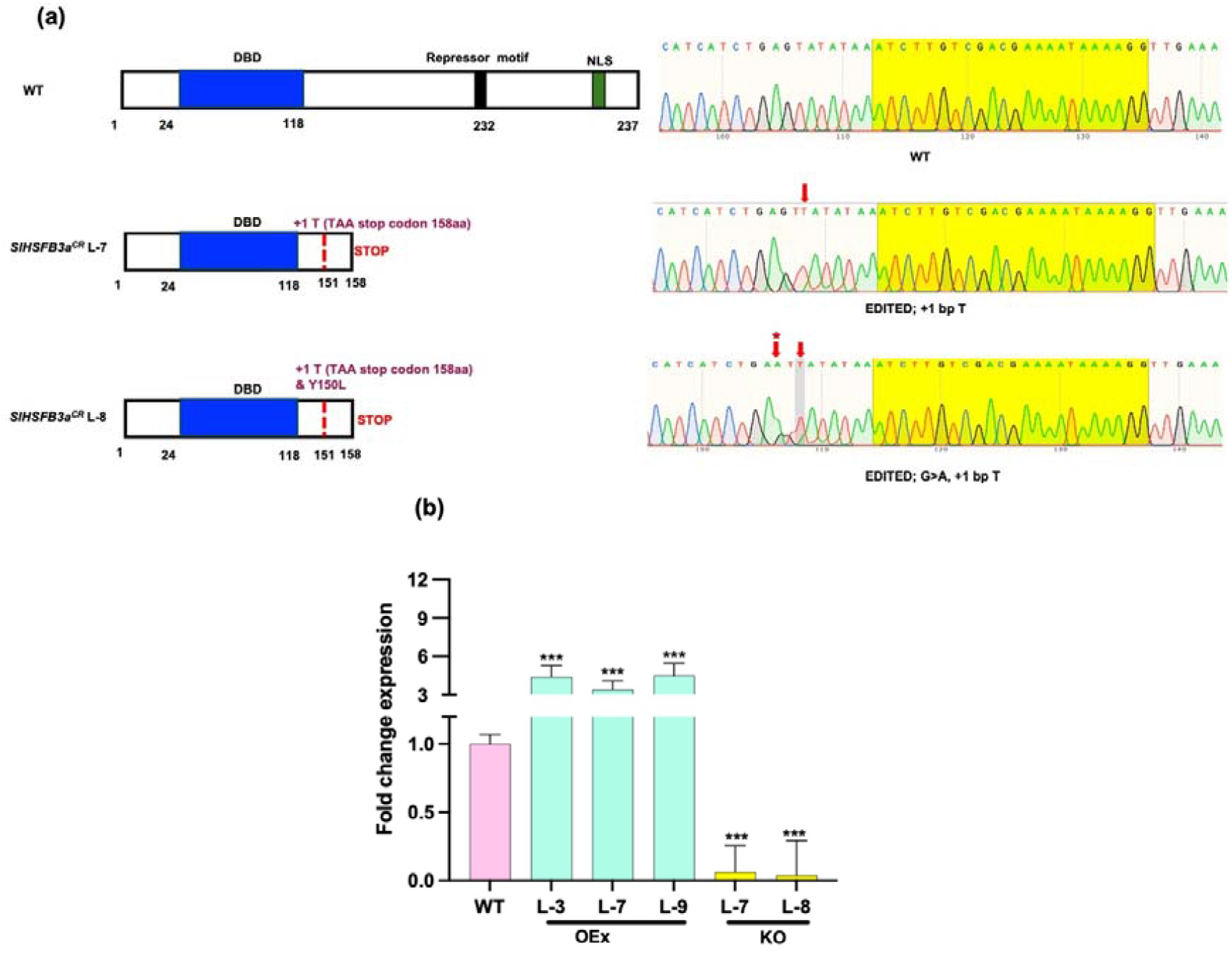
Sanger sequencing of *SlHSFB3a* CRISPR-edited lines and *SlHSFB3a* transcript validation in OEx and KO. (A) Gene structure and Sanger sequencing of the edited gene fragment for mutation detection, along with WT *SlHSFB3a* genomic sequence. GuideRNA along with PAM site is yellow color-coded. Red arrows denote indels. (B) *SlHSFB3a* transcript quantification in OEx and KO via reverse transcription quantitative polymerase chain reaction, normalized against *SlCAC*. Error bars represent the standard error (±) calculated from three independent biological replicates. Significant variation is indicated by asterisks *P < 0.05, **P < 0.01, and ***P <0.001 (single factor ANOVA).

**Fig. 3:**
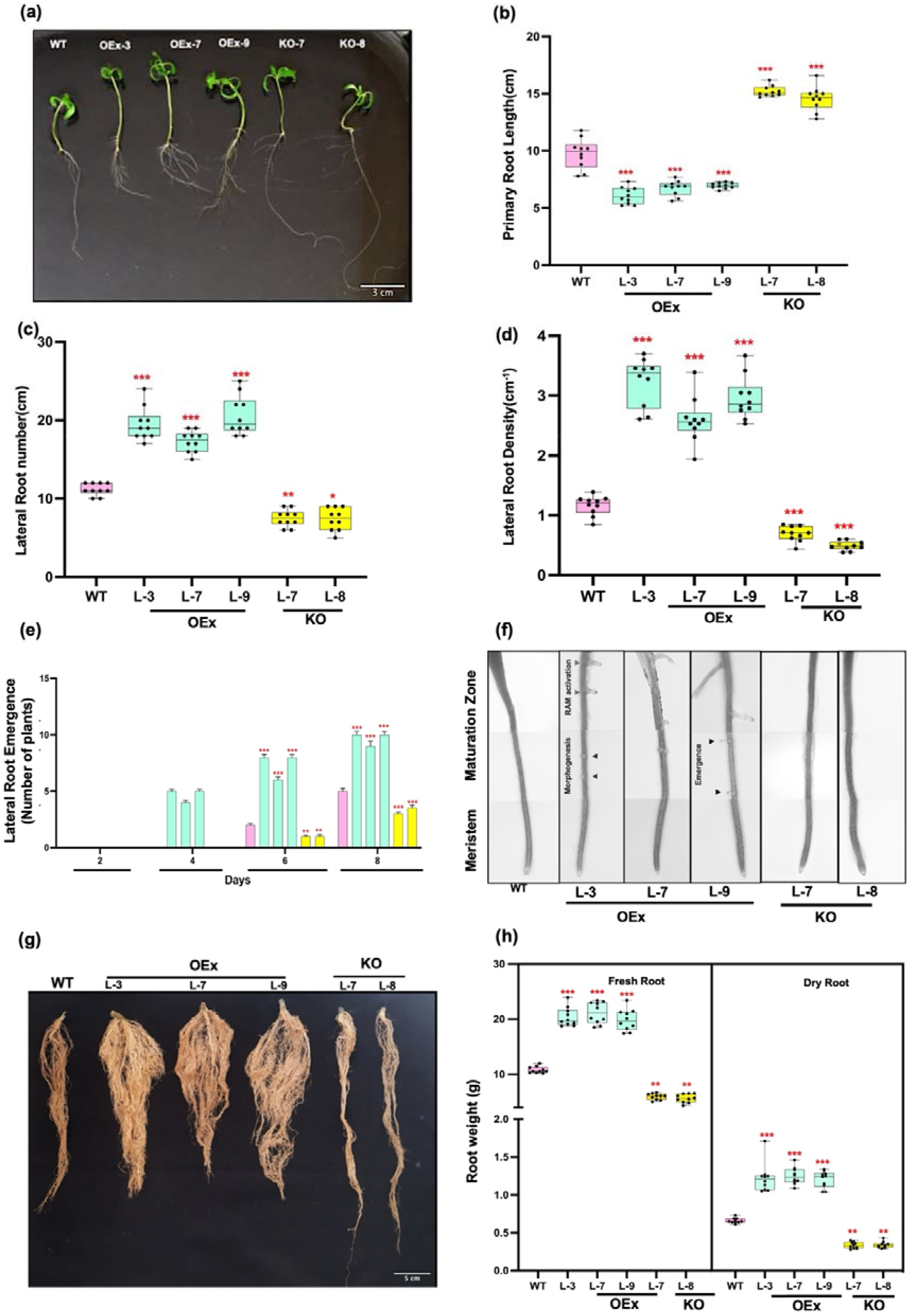
Effect of *SlHSFB3a* manipulation in tomato root phenotype. (A) Visual image of 15DAG seedlings of *SlHSFB3a* OEx, KO and WT. Quantification of root parameters (B) Primary root length (cm) (C) Lateral root number (D) LR Density (E) LR emergence and stereomicroscopic images of 4DAG seedlings demonstrating early LR emergence in OEx (F). (G) and (H) Root weight (g) of mature grown *SlHSFB3a* transgenics (60 DAG), under normal conditions. Represented data are the averages, and standard error is from three independent experiments. Data points indicate each biological replicates.

### *SlHSFB3a* enhances root biomass and volume in mature tomato plants

*SlHSFB3a* levels significantly affected root phenotypes in seedlings (15 DAG) under *in vitro* conditions. However, *SlHSFB3a* displays highest expression in 60 DAG roots, and this guided the focus towards observing root phenotypic changes during later or mature plant stage. In OEx, both fresh and dry root weights increased significantly by 84-96%, while KO showed 45-48% reduction compared to WT (Fig. **3g, h**). Also, root volume measurements revealed a substantial increase in OEx roots, with KO roots displaying only one-fifth of the WT root volume (Supplementary **Fig. 3**). These results suggest that *SlHSFB3a* overexpression enhances root branching, contributing to increased root biomass and volume in mature, soil-grown plants, underscoring its role in root development.

### *SlHSFB3a* role in regulating signaling networks in tomato root growth

To elucidate the transcriptional regulatory mechanisms of *SlHSFB3a*, RNA sequencing of WT, OEx, and KO roots (30 DAG) grown in greenhouse was carried out (Fig. **4a**). Deep RNA sequencing yielded 62.2 to 103.2 million total raw reads, with high-quality reads aligning against the tomato reference genome (*Solanum lycopersicum* SL3.0.54) (Supplementary Table 2). Volcano plots and heatmaps indicated that genes upregulated in KO were statistically significant, correlating with the observed expression patterns (Fig. **4b, d**). Conversely, in OEx, downregulated genes were more significant. Differentially expressed genes (DEGs) analysis revealed changes in genes involved in auxin signaling pathways (Fig. **4 e, f**; Supplementary Table 3). Particularly auxin repressors *SlIAA2* and *SlIAA3* expression showed significant changes in OEx and KO (Fig. **4h**). *SlIAA3* was suppressed in OEx lines and upregulated in KO, while *SlIAA2* displayed similar contrasting patterns, suggesting *SlHSFB3a’s* repressor role in auxin-mediated root architecture regulation (Supplementary Table 3). *SlARF6b* and *SlLOB42*, two LR related genes showed higher expression in OEx and repressed/undetermined in KO. The transcript levels of auxin transporters such as *SlPIN7, SlPIN8*, and *SlLAX5*, were enhanced in OEx but remained undetectable in KO under experimental conditions, suggesting *SlHSFB3a’s* influence on auxin biosynthesis and signaling in roots (Fig. **4h**, Supplementary Table 3).

**Fig. 4:**
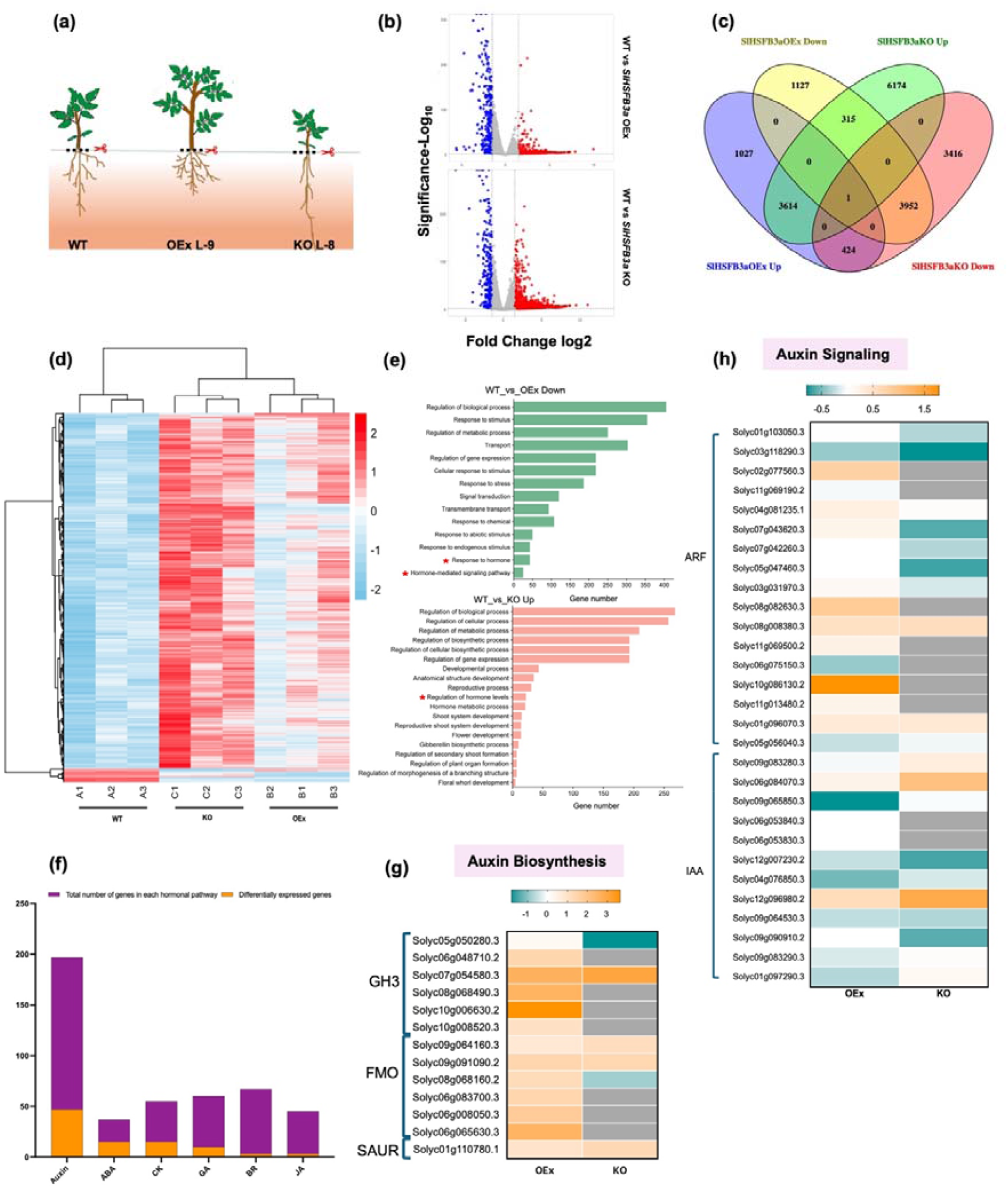
Transcriptomic responses of *SlHSFB3a* manipulation in tomato roots (A) RNA seq strategy, and (B) Volcano plots showing up and down DEGs proportion in two combinations, WT vs *SlHSFB3a*OEx and WT vs *SlHSFB3a*KO. (C) Venn diagram quantifying common and unique DEGs in WT and *SlHSFB3a* OEx_Up, *SlHSFB3a* OEx_Down, *SlHSFB3a* KO_Up and *SlHSFB3a* KO_Down. (D) Heatmap showing most DEGs upregulated in KO as compared to OEx and WT. (E) Functional annotation of DEGs in two combinations using *Solanum lycopersicum* reference genome in The Gene Ontology Resource. (F) Hormone related DEGs in RNA seq data. (G) and (H) DEGs significantly expressed related to auxin biosynthesis and signaling.

### *SlHSFB3a* modulates auxin sensitivity in tomato roots

RNA sequencing analysis showed that most DEGs in *SlHSFB3a* transgenic roots were linked to auxin biosynthesis and signaling. To study the effect of exogenous auxin on LRs in *SlHSFB3a* manipulated seedlings, a dose-dependent IAA treatment was applied over 10 days (Fig. **5a**). LR number increased significantly in a dose dependent manner in WT upon exogenous IAA treatment. WT LRs showed significant increase from 6.8 ± 0.4 (mock) to 14.5 ± 4 upon 10 nM IAA treatment. At low IAA (10 nM), OEx plants showed minimal increase in LR count 3.57%, 9.45% and 11.03% in three individual lines as compared to WT. However, at higher IAA concentrations (50 nM and 100 nM) OEx LR quantity increased ranging from 77.92 to 103.37%, upon comparison with their respective WT roots (Fig. **5b**). KO plants, however, responded more readily to 10 nM and 50nM IAA but showed reduced LR numbers at 100 nM, confirming *SlHSFB3a*’s influence on auxin sensitivity.

**Fig. 5:**
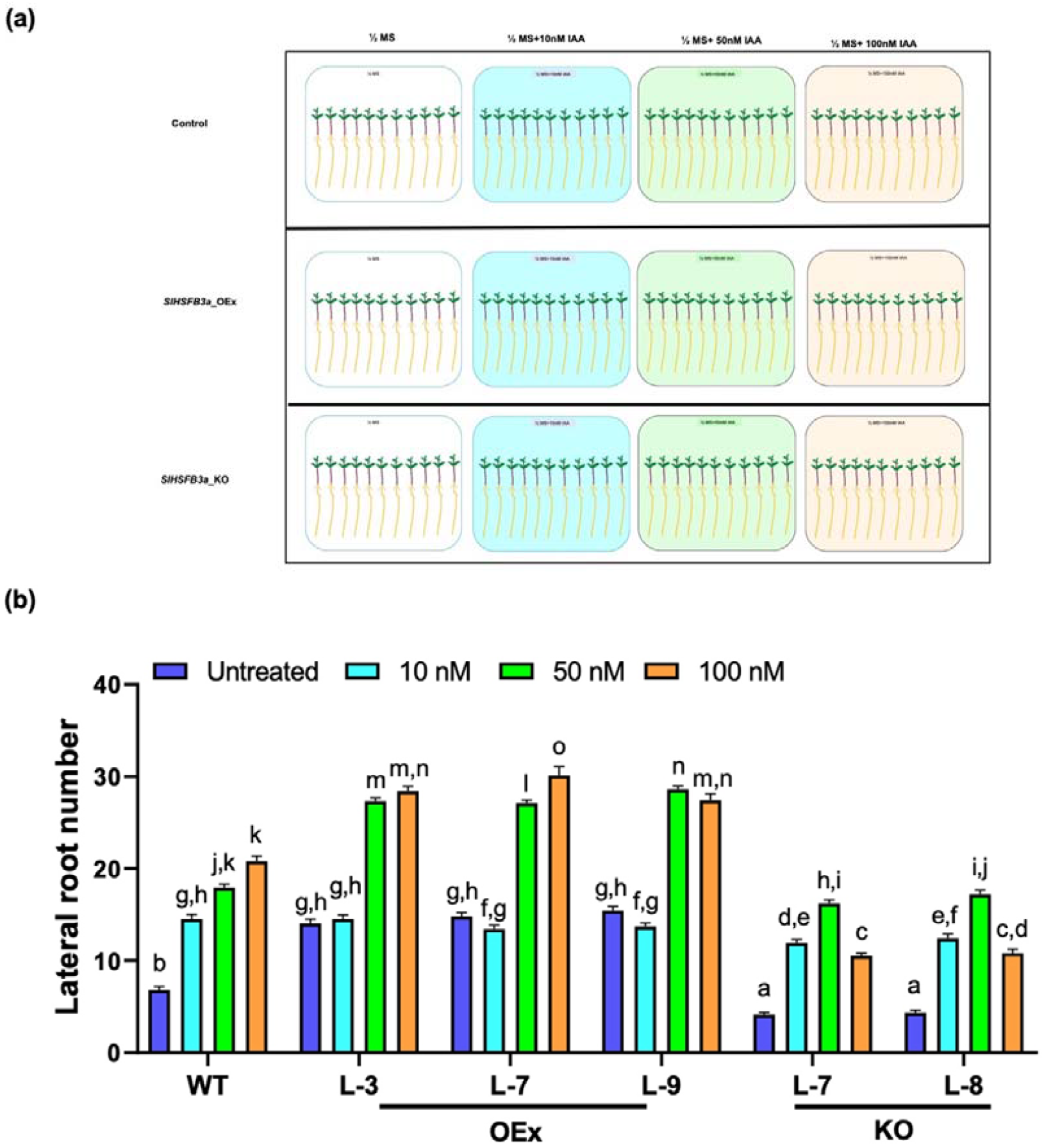
Exogeneous IAA treatment in OEx and KO tomato seedlings. (A) Experimental setup; From top to bottom WT, OEx and KO seedlings treated according to increasing IAA concentrations and untreated (left to right). (B) LR quantification of IAA treated OEx, KO and WT (15 DAG). Alphabets denotes significant differences across concentration gradient treatment groups (One-way ANOVA, DMRT *post-hoc*).

### *SlHSFB3a* represses auxin signaling repressors and promotes LR growth

*SlIAA2* and *SlIAA3*, known auxin repressors, are differentially expressed in OEx and KO (Fig. **4h**, Supplementary Table 3). *SlHSFB3a*, being a transcriptional repressor in nature, likely targets *SlIAA2* and *SlIAA3* directly to modulate auxin sensitivity, thereby promoting lateral root initiation and growth. RT-qPCR validated these RNA-seq findings, showing significant reduction in *SlIAA3* and *SlIAA2* expression in OEx (Fig. 6**a, b**). *SlIAA2* transcripts were slightly elevated in KO but significantly repressed by 54-65% in three OEx transgenics. Although greenhouse grown mature KO plants (30 DAG) showed much increase in *SlIAA2* transcripts ~4 fold. Faint *SlIAA3* expression in OEx but significantly elevated in two individual KO lines around ~23 and ~58 fold as compared to WT (Fig. **6a, b**).

**Fig. 6:**
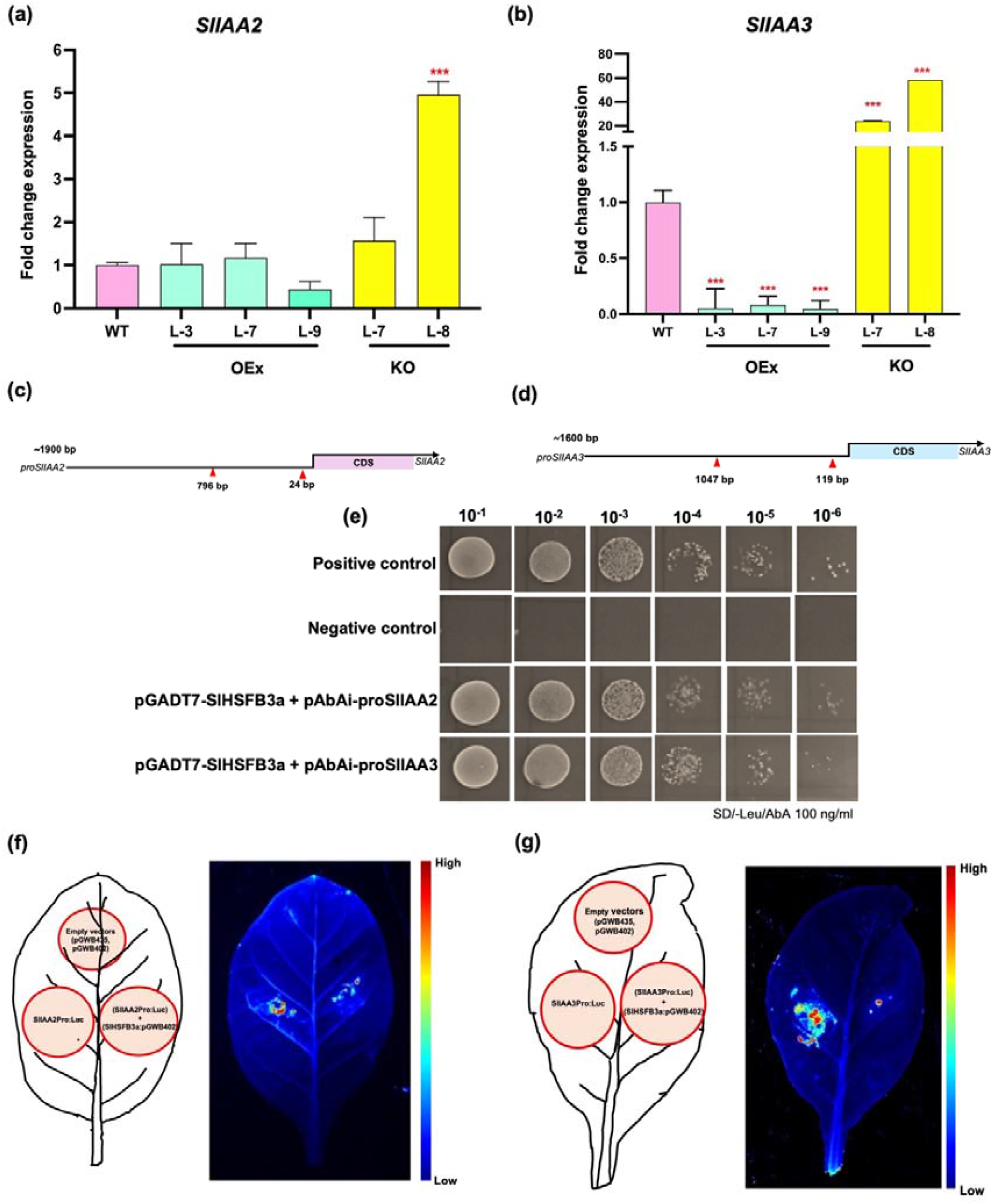
*SlHSFB3a* directly regulates the expression of *SlIAA2* and *SlIAA3*, by directly binding to their promoters. RT-PCR of (A) *SlIAA2* and (B) *SlIAA3* expression in OEx, KO and WT. *SlCAC* is used as housekeeping gene. Schematic diagrams of (C) *SlIAA2* and (D) *SlIAA3* promoters highlighting the HSE motif location. Intense interaction between *SlHSFB3a* and *SlIAA2* and *SlIAA3* promoters confirmed via (E) yeast one hybrid assay; (F) and (G) Dualluciferase assay. The reduced effect of luciferase assay confirms direct interaction due to SlHSFB3a repressor nature. Three independent replicates were considered.

*In silico* analysis identified HSE (Heat Stress Element) cis-acting motifs in the *SlIAA2* (+374 bp, +417 bp) and *SlIAA3* (+1095 bp, +1867 bp, +2003 bp) promoters, indicating potential binding sites for *SlHSFB3a* (Fig. 6**c, d**). Yeast one hybrid confirmed strong interaction between *SlHSFB3a* and *SlIAAs* (Fig. 6**e**). To compare the expression pattern between *SlHSFB3a* and *SlIAA* promoters *in-planta*, dual-luciferase assay was done. Co-infiltration of pGWB402:*SlHSFB3a* along with pGWB435:*SlIAA2Pro* and pGWB435:*SlIAA3Pro* in *Nicotiana tabacum* leaves resulted in reduction of promoter activity (Fig. 6**f, g**), thereby clearly demonstrating that *SlHSFB3a* represses *SlIAA2* and *SlIAA3* genes *in planta* by directly binding to their promoter regions. Altogether, these findings suggest that *SlHSFB3a* negatively regulate auxin signaling repressors *SlIAA2* and *SlIAA3*, which directly alter auxin signaling and thereby promote lateral root growth in tomato roots.

By correlating the observed root phenotype with the transcriptomics data, few other components of the auxin signaling pathway that are root-related were identified. *SlARF7* and *SlARF19*, activators of auxin signaling were upregulated in *SlHSFB3a* OEx (Fig. **7a, b**). *SlARF7* showed ~129 folds, ~26 folds and ~17-fold increase in OEx, and was undetectable in KO, this suggests that *SlHSFB3a* positively regulates *SlARF7* by regulating the expression of *SlIAA2* and *SlIAA3* (Fig. **7a**). LOBs, or lateral organ boundary domain transcription factors, are known to play roles in lateral organ initiation, organ patterning, root growth, and plant regeneration. The increased expression of *SlLOB42* and *SlLOB20* (Fig. **7c**; Supplementary Table 3) in OEx suggests that these genes might be controlled via *SlIAA2* and *SlIAA3* signaling cascade acts downstream of *SlHSFB3a*, potentially contributing to the regulation of lateral root initiation in tomato.

**Fig. 7:**
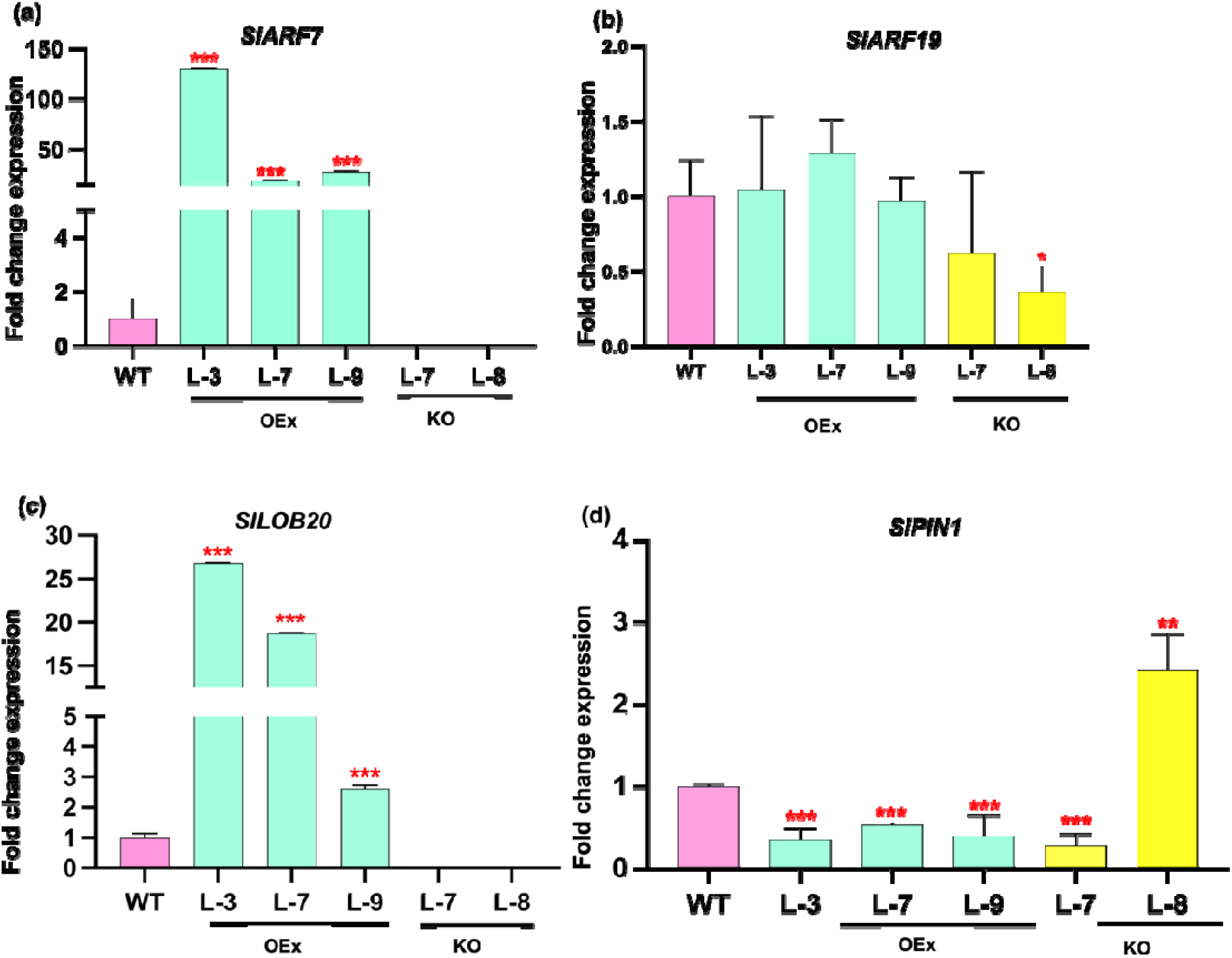
*SlHSF3a* modulates root-related auxin response genes. Graphical representation of (A)*SlARF7*, (B) *SlARF19*, (C) *SlPIN*, and (D) *SlLOB20* expression in OEx KO and WT. Transcript levels were normalized against *SlCAC*. Y-axis depicts 2^−ddCT^. Bar graph represents the mean values with standard error from three biological replicates. Significant variation is represented by asterisks *P < 0.05, **P < 0.01, and ***P <0.001 (one-way ANOVA).

## Discussion

A thorough comprehension of intricate genetic nexus determines root architecture that is a prerequisite for breeding improved crop varieties. Numerous studies have explored root patterning in plants, it is important to note that root structures differ among species depending on their geographical distribution and adaptive behaviors. Hence, this study was undertaken to focus on the genetic and hormonal control behind dicot root development. Previously, a tomato tissue-specific transcriptomics (Kumar *et al*., 2021) identified several root-specific genes, including TFs such as two members of the HSF family, that showed predominant expression in roots compared to other tissues. Plant HSFs are one of the early transcription factor families that have adapted since the start of terrestrial evolution. Their function as “master thermoregulators” evolved through genetic and molecular changes in response to extreme environment. HSFA first appeared during the chlorophyta stage, later expanding to HSFB and HSFC. Functional divergence within the HSF family occurred as angiosperms evolved into monocots and dicots (Wang *et al*., 2018). In tomato, the expansion of the HSF gene family resulted from segmental and tandem gene duplications (Majee *et al*., 2023). Notably, *SlHSFB3a* and *SlHSFB5* were identified only in dicots, highlighting their potential role in dicot-specific development (Majee *et al*., 2023).

*SlHSFB3a*, a class B HSF, has not yet been characterized. Its specific expression in age-dependent tomato root tissues suggests its involvement in root growth regulation. Present study functionally characterizes *SlHSFB3a*, and its association with root development under normal/unstressed conditions. This association was unveiled through a comprehensive approach that includes expression profiling, sequence analysis, and it’s functional characterization in tomato by generating genetically manipulated transgenics. *SlHSFB3a*’s in involvement in tomato root growth was marked upon by its high expression in age-dependent roots that induces upon ABA, IAA, and BAP, and suppresses by JA, SA, and ACC {Fig. **1(d-i**)} treatment. These findings align with previous reports linking HSF expression to various hormones (Fragkostefanakis *et al*., 2015; Hu *et al*., 2015; Huang *et al*., 2015; Zhang *et al*., 2015). In Tea, *CsHSFA3, CsHSFB1*, and *CsHSFC1* were upregulated by ABA, while rest were downregulated (Zhang *et al*., 2020). Muthuramalingam et al. (2019)reported the transcript profiling of 25 rice HSFs were upregulated by ABA and JA and downregulated by GA, cytokinin, and auxin (Muthuramalingam *et al*., 2020).

Additionally, phenotypic analysis of genetically modified *SlHSFB3a* transgenics improved root architecture under both *in-vitro* and greenhouse conditions. The contrasting phenotypes of OEx and KO clearly demonstrate that while *SlHSFB3a* suppresses primary root growth, it positively regulates LR growth (Fig. **3b, c**), enhancing LRD. LR formation involves initiation, emergence, and outgrowth (Du & Scheres, 2018). *SlHSFB3a* promotes early LR initiation and emergence. Likewise, improved root morphology was reported in Begum et al. (2013) by overexpression of *AtHSFB4* in Arabidopsis (Begum, Reuter & Schöffl, 2013). Under greenhouse conditions, a key finding is enhanced root volume in *SlHSFB3a* OEx roots. A single root spans a specific volume, extending from the initial root zone to the tip of the main root, with lateral roots (LRs) branching out and spreading through the surrounding soil (Hinsinger *et al*., 2005). Roots occupy 2-3% of the topsoil, root volume plays a vital role in determining the depth of soil penetration. Increased root volume improves crop quality by facilitating better oxygenation in plants (Redillas *et al*., 2012; Xu *et al*., 2013; Hoffmann *et al*., 2020), offering significant agro-economic advantages. Mature *SlHSFB3a* OEx plants exhibited higher root volume compared to WT plants, whereas KO plants showed a marked reduction of 80-81% in root volume (Supplementary Fig. **3**).

Data mining of RNA seq data revealed majority of DEGs related to hormone signaling and growth. Amongst them, DEGs involved in auxin signaling AUX/IAAs, ARFs, LOBs, and few auxin transporters like *SlPIN7, SlPIN8*, and *SlLAX5* (Fig. **4h**, Supplementary Table 3) showed significant expression. We identified two auxin repressors, *SlIAA2* and *SlIAA3* to be expressed in KO. *SlIAA3* showed its expression is restricted to root caps and LRP in tomato (Chaabouni *et al*., 2009). In *Arabidopsis*, the *SHY/IAA3* complex functions in the endodermis of the differentiation/maturation zone and is induced by auxin released by the emerging LRP. *SlARF6b* and *SlARF7* were induced in OEx and showed contrasting levels in KO roots. This co-relates with several studies in *Arabidopsis* and other crops, ARF7 and ARF19 are crucial in LR initiation and patterning. This activation drives downstream auxin signaling, including the induction of genes like *SlLOB20* and *SlLOB42* (supported by RNA seq and qRT-PCR data), that might facilitate LR emergence and increase LRD. LOB20 is not much characterized, but few studies suggest their root-specific role. In cassava, *MeASLBD20* expression in roots are significantly high as compared to other tissues (Mao *et al*., 2023). LOB20 is undetectable in shoots, but abundant in *Arabidopsis* roots (Thatcher *et al*., 2012), otherwise demonstrated to be involved in pathogen defense. *SlPIN7* and *SlPIN8*, key auxin transporters, were upregulated in OEx and undetected in KO, indicating their role in maintaining auxin homeostasis. *AtPIN8* regulates LR emergence (Cui *et al*., 2022), and its loss-of-function mutation results in reduced LR density, a phenotype similar to that observed in *SlHSFB3a* KO. The increased expression of auxin transporters like *SlPIN7* in OEx further supports the idea that *SlHSFB3a* regulates tomato root physiology by maintaining auxin concentrations. These outcomes agree with previously reports (De Rybel *et al*., 2010; Ruiz Rosquete, Waidmann & Kleine-Vehn, 2018) showing that lateral root growth involves auxin signaling, but the novel outcome is that an HSF submember transcriptional regulation is a part of it.

Downregulation of *SlIAA2* and *SlIAA3* in OEx, contrasting in KO (validated by RT-PCR and RNA seq), are responsible for increased auxin signaling and LR number. Y1H and dual luciferase confirmed direct interaction between *SlHSFB3a* and IAA promoters. The novelty of this study lies in the fact that the recruitment of HSFB3a to downstream *SlIAA* promoters is directly linked to it’s transrepression activity.

Our experiments form part of a broader initiative to identify key genetic factors influencing root growth in tomato. This study identifies *SlHSFB3a* as a pivotal regulator of tomato root development, primarily through its modulation of auxin signaling. By suppressing auxin inhibitors, *SlHSFB3a* enhances lateral root formation and overall root growth, establishing its role as a central determinant of tomato root system (Fig. **8**). Moreover, our study highlights the importance of HSF transcription factor submember in plant growth under normal/unstressed conditions, shedding light on their previously underexplored biological function. As a novel path has been unraveled for HSFs (previously tagged as “thermoregulators”) future research will delve into in-depth interactions between *SlHSFB3a* and other genetic factors such as *SlLOB20, SlLOB42* and decode their expression pattern root signaling. These findings offer promising avenues for the development of improved tomato varieties by employing these genetic factors as markers. Given the critical role of optimized root architecture in promoting plant vigor and productivity, these findings hold considerable potential for agrocommercial applications.

**Fig. 8:**
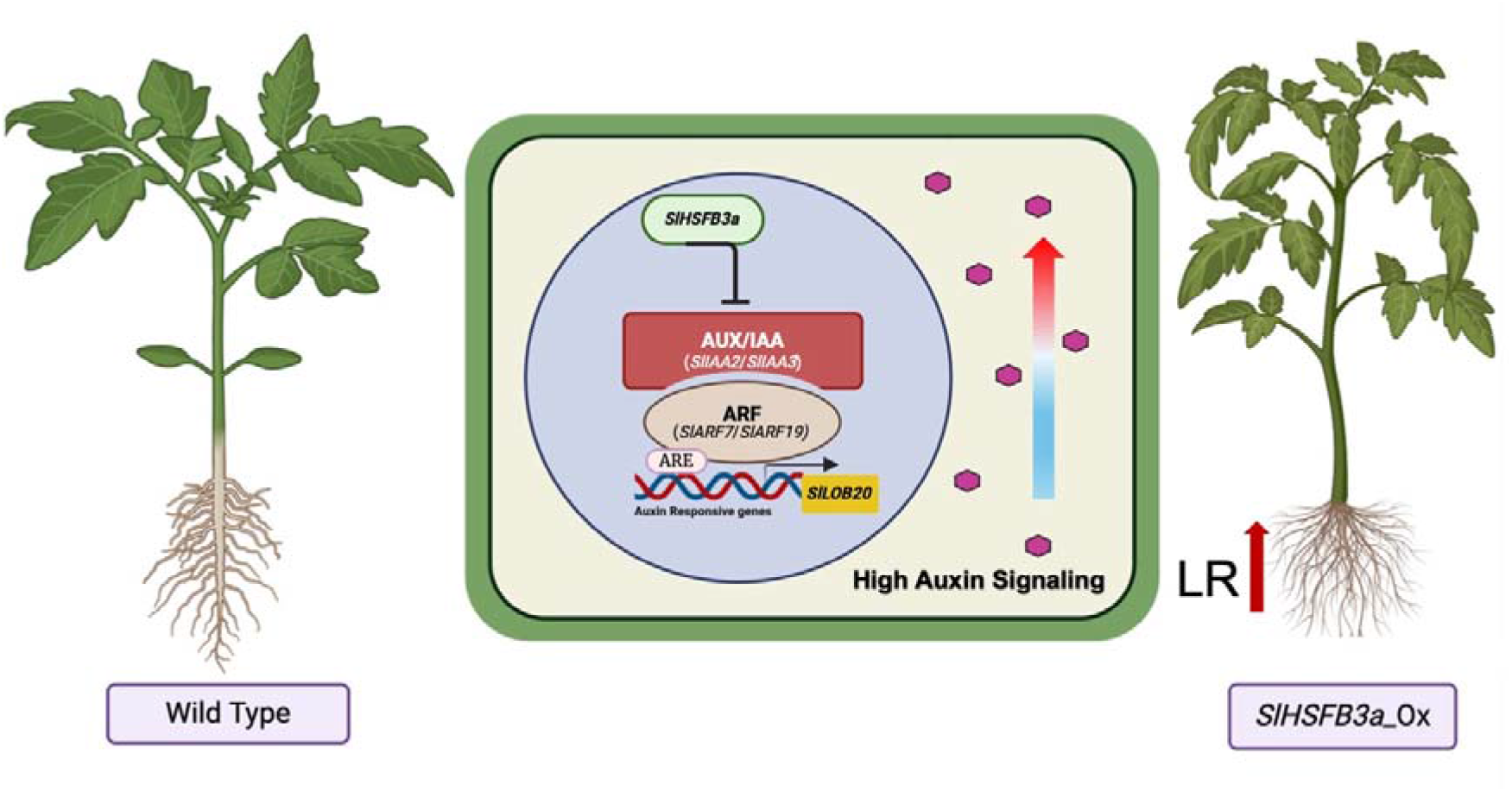
Proposed mode of interaction between *SlHSFB3a* and key auxin signaling components regulating LR growth and enhancing tomato root architecture. Under normal conditions, *SlHSFB3a* emerged as a central participant in regulating LR growth in tomato via auxin pathway. *SlHSFB3a* being a transrepressor in nature, suppressing auxin repressors *SlIAA2* and *SlIAA3* as direct target genes. This leads to high auxin signaling inside the root cell. With low expressed AUX/IAAs, *SlARF7* and *SlARF19* is upregulated. Higher transcript abundance of *SlARFs*, activates the transcription of auxin responsive gene *SlLOB20* by interacting via ARE motifs present in the promoter. This enhances LR number and density in *SlHSFB3a* OEx tomato roots.

## Conclusion

Our findings reveal that *SlHSFB3a* serves as a pivotal regulator of root growth in tomato by orchestrating genetic control through its temporal expression and auxin-mediated downstream signaling pathways. Under normal, unstressed conditions, *SlHSFB3a* fine-tunes the equilibrium among auxin signaling components, acting as a key modulator of auxin sensitivity. This regulatory mechanism involves direct modulation of key auxin repressors, *SlIAA2* and *SlIAA3* by *SlHSFB3a*, to maintain auxin homeostasis in root meristem. Repressed AUX/IAAs increase auxin signaling activating *SlARF7* and *SlARF19* that are responsible LR development. Taken together, *SlHSFB3a* fine-tunes the balance between auxin signaling components and acts as a vital modulator of auxin sensitivity and affects root architecture

## Acknowledgments

We thank Dr Shucai Wang (Northeast Normal University, Changchum China) for the effector and reporter constructs for the repressor assay and Dr. Prabodh K. Trivedi, NBRI, Lucknow for sharing pHSE401 vector. Ram Awadh for taking care of plants in greenhouse. AM is financially supported by DST-INSPIRE as a Senior Research Fellow, SBT and VK received SRF funding from CSIR, India.

## Competing Interest None declared Author contributions

VAS and AAPS conceptualized and conceived the idea. AM, BS and VK designed the experimental pipeline and conducted them. AM and VK contributed to statistical and data analysis. AM and VAS drafted the first manuscript version. All authors provided their input for the final manuscript.

## Data Availability

The raw data of the RNA seq analysis have been submitted to the SRA data under the Bioproject Accesssion number PRJNA1227233.

