## Supplementary figures and images for "*SlHSFB3a* developmentally regulates lateral root formation by modulating auxin signaling in tomato"

### Supplementary Figures S1-S4

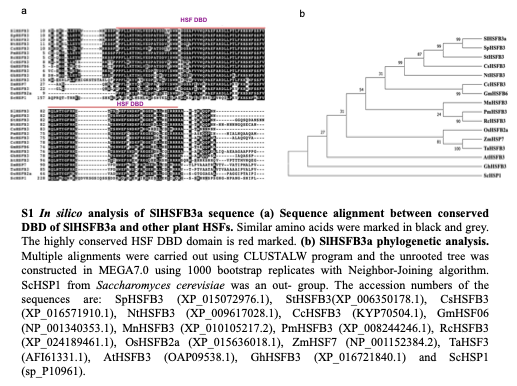

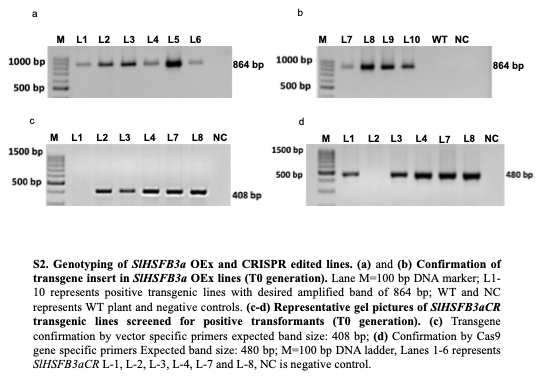

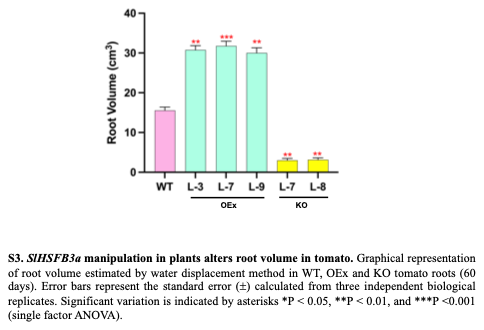

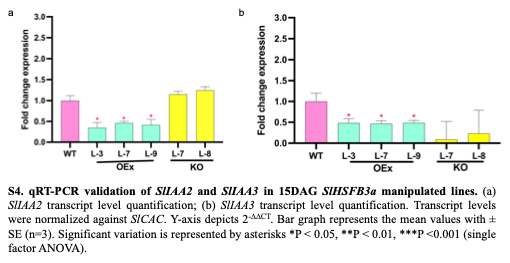
